# Gpr125 identifies myoepithelial progenitors at tips of lacrimal ducts and is essential for tear film

**DOI:** 10.1101/2020.09.15.296749

**Authors:** Elena Spina, Rebecca Handlin, Julia Simundza, Angela Incassati, Muneeb Faiq, Anoop Sainulabdeen, Kevin C Chan, Pamela Cowin

**Affiliations:** Departments of Cell Biology, New York University School of Medicine; Departments of Dermatology, New York University School of Medicine; Departments of Ophthalmology, New York University School of Medicine

## Abstract

Gpr125, encoded by *Adgra3*, is an orphan adhesion G-protein coupled receptor (aGPCR) implicated in modulating Wnt signaling and planar polarity. Here we establish both physiological and pathological roles for Gpr125. We show that mice lacking Gpr125 or its signaling domains display an ocular phenotype with many hallmarks of human dry eye syndrome. These include squinting, abnormal lacrimation, mucus accumulation, swollen eyelids and inflammatory infiltration of lacrimal and meibomian glands. Utilizing a Gpr125-β-gal reporter and scRNAseq, we identify Gpr125 expression in a discrete population of cells located at the tips of migrating embryonic lacrimal ducts. By lineage tracing we show these cells function as progenitors of the adult lacrimal myoepithelium. Beyond defining an essential role for Gpr125 in tear film and identifying its utility as a marker of lacrimal progenitors, this study implicates Gpr125 in the etiology of blepharitis and dry eye syndrome, and defines novel animal models of these common maladies.

## Introduction

Gpr125 is an orphan adhesion G-protein coupled receptor (aGPCR) that was discovered through homology searches of the human genome database (Bjarnadottir et al., 2006). Like other members of this family, Gpr125 has a large extracellular domain with sequence similarity to cell adhesion molecules (Figure 1A) (Bjarnadottir et al., 2006; Simundza and Cowin, 2013). Previous studies have highlighted Gpr125 as a marker of undifferentiated murine spermatogonial progenitors (Seandel et al., 2007), documented its elevation in the choroid plexus following injury and correlated high Gpr125 expression with both good and poor outcome in cancer (Fu et al., 2013; Pickering et al., 2008; Wu et al., 2018). When introduced into cultured cells Gpr125 undergoes constitutive clathrin-mediated internalization to endosomes suggesting a role in receptor recycling (Spiess et al., 2019). When expressed ectopically in zebrafish, Gpr125 interacts with the cytoplasmic adaptor Disheveled (Dsh) and recruits Frz7 and Glypican4 (Gpc4) complexes. When reduced it has little effect in wildtype zebrafish but in Wnt/planar cell polarity (PCP) mutants exacerbates defects in convergent extension and directed migration of facial branchiomotor neurons (FBMN) (Li et al., 2013). Gpr125 shares significant homology with Gpr124, which has been shown to regulate angiogenic sprouting and control selective Wnt signaling by stabilizing specific ligand receptor interactions (Anderson et al., 2011; Vanhollebeke et al., 2015). To date, the physiological function of endogenous Gpr125 in higher vertebrates has remained elusive. Here we uncover an essential physiological role for Gpr125 in the lacrimal gland, and a pathological role in the etiology of blepharitis and dry eye disease (DED).

**Figure 1.**
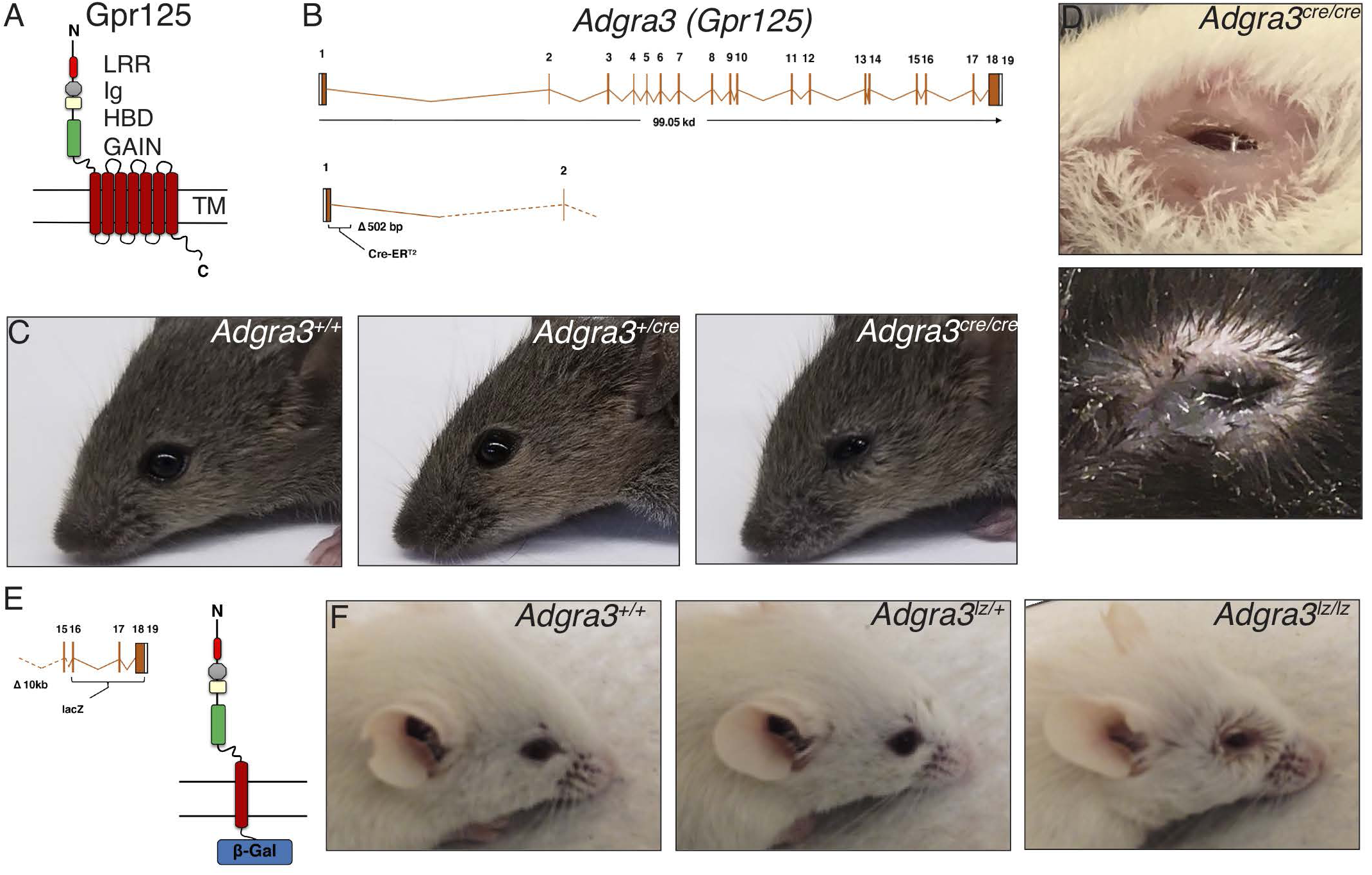
Gpr125 loss induces blepharedema, blepharitis and mucus accumulation. **(A)** Schematic of Gpr125 protein comprising N-terminus (N), leucine rich repeats (LRR), Immunoglobulin-like domain (Ig), hormone binding domain (HBD), GPCR autoproteolyis-inducing (GAIN) domain, 7-pass transmembrane region (TM) and cytoplasmic region (C). (**B)** Schematic of *Adgra3*. *Adgra3^cre/cre^* mice were generated by replacement of 502bp after the first codon with a creER^*T2*^ module. (**C)** Eye phenotype of *Adgra3^cre/cre^* compared to controls. **(D)** Examples of blepharedema and mucus accumulation in *Adgra3^cre/cre^* mice. **(E)** Schematic of the Gpr125-β-gal protein generated by deletion of 10 kb sequence downstream of the first TM and replacement by *lacZ*. **(F)** Eye phenotype of *Adgra3^lz/lz^* mice compared to controls.

## Combined Results & Discussion

### Mice lacking Gpr125 display blepharitis, blepharedema and mucoid accumulation

To address the role of native Gpr125, we developed mice that permit Gpr125 expression to be ablated and the lineage of cells normally expressing it to be traced by inserting a creER^*T2*^ cassette downstream of the *Adgra3* promoter (Figure 1B). Mice lacking Gpr125 expression (*Adgra3^cre/cre^*) display a prominent eye phenotype (Figure 1C). *Adgra3^cre/cre^* mice squint as soon as their eyes open; whereas, heterozygous *Adgra3cre/+* are indistinguishable from wild-type littermates (Figure 1C). As *Adgra3^cre/cre^* mice mature, this early blepharitis progresses to blepharedema (swollen balding eyelids) and mucus precipitation (Figure 1D). The phenotype is constant in males but in females oscillates with reproductive status, becoming pronounced during pregnancy and lactation. During these stages, mice develop proptosis (bulging eyes) that resolves during weaning. The eye phenotype in *Adgra3^cre/cre^* mice is 100% penetrant on all strain backgrounds examined (C57B6/CH3, FVBN, and mixed). To dissect the role of Gpr125’s adhesion ectodomain from its internal signaling functions we examined a second strain, *Adgra3^lz/lz^*, which expresses the Gpr125 ectodomain and 1^st^ transmembrane domain fused in frame to β-galactosidase and lacks regions required for signaling/adaptor functions (Figure 1E) (Seandel et al., 2007). Homozygous *Adgra3^lz/lz^* mice recapitulate the *Adgra3^cre/cre^* null phenotype (Figure 1F), whereas, *Adgra3^lz/+^* mice are normal. Collectively, these data demonstrate that Gpr125 has an essential physiological role in normal eye development and indicate that signaling downstream of the receptor is required. These analyses also reveal that loss of Gpr125 protein or Gpr125 signaling is sufficient to trigger several common eye pathologies, such as blepharitis, blepharedema and mucoid accumulation.

### Gpr125 in ocular structures and correlated pathologies

We examined adult eye globes by X-gal staining and in *Adgra3^lz/+^* mice found strong Gpr125-β-gal expression in the inner layer of the iris and in the ciliary body, which secretes aqueous humor (Figure 2A, B). As abnormal aqueous humor dynamics can alter intraocular pressure (IOP), which is a major risk factor for glaucoma, we measured IOP, but found no significant difference between wildtype and mutant genotypes (Figure 2C).

**Figure 2.**
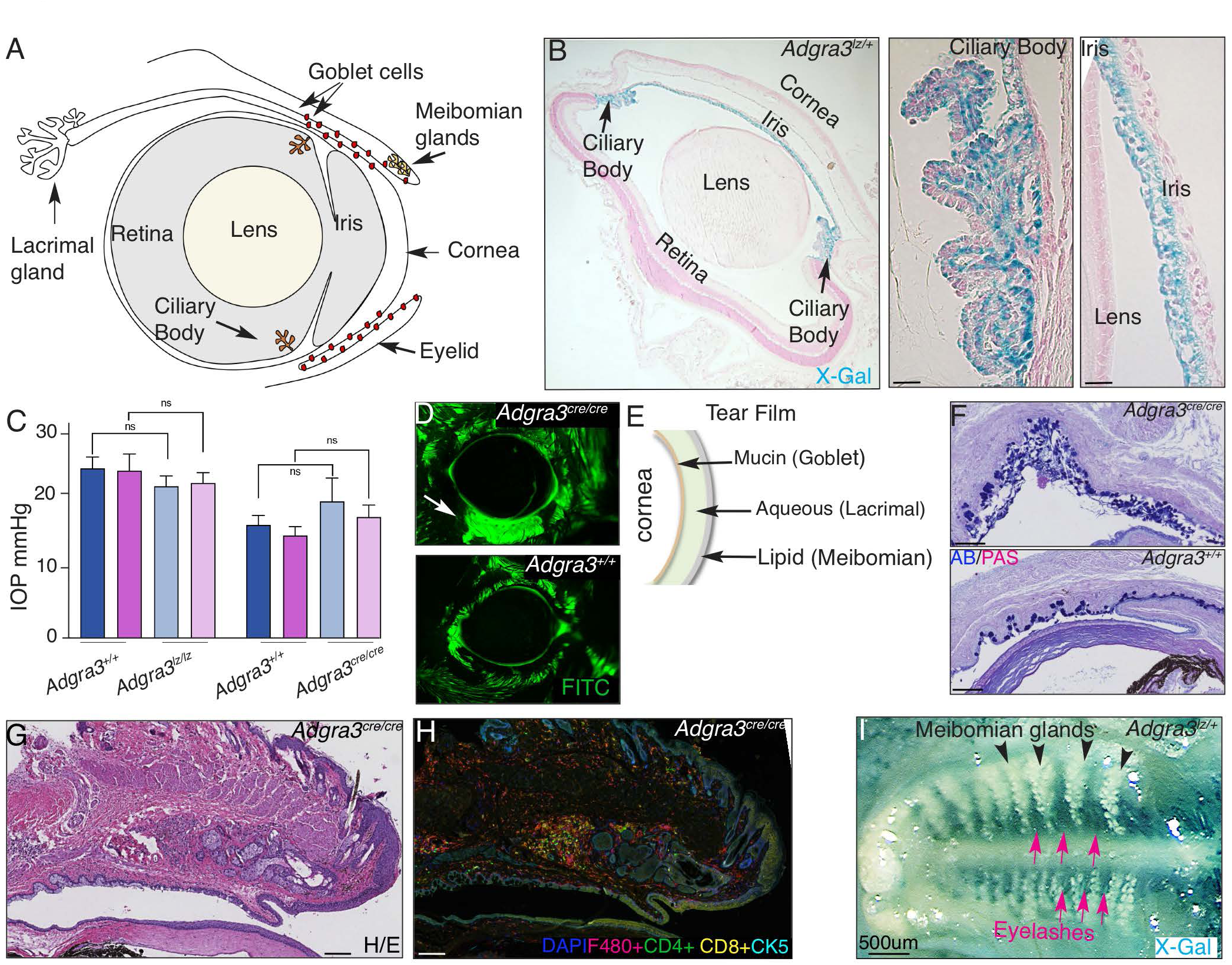
Gpr125 is expressed in eyes and eyelids. **(A)** Diagram of murine eye. (**B)** Section of X-gal stained *Adgra3^lz/lz^* eye shows Gpr125-β-gal expression in the ciliary body and iris. (**C)** Intraocular pressure (IOP) (mmHg) in male (blue) and female (pink) *Adgra3^lz/lz^* and *Adgra3^cre/cre^* mice compared to their respective FVBN and B6 controls. Each bar represents the mean ± SEM on 6-14 mice/group. ns, not significant. (**D)** Fluorescein stained corneas in *Adgra3^cre/cre^* and control mice. n=3 (**E)** Schematic of tear film. (**F)** Eyelid sections stained with alcian blue AB/PAS show goblet cells in *Adgra3^cre/cre^* and control mice. n=23. **(G-H)** Sections of eyelids showing meibomian glands stained with **(G)** H/E and **(H)** immunostained with antibodies: F480, CD4,CD8, CK5 and DAPI to detect macrophages, T-helper, cytotoxic T cells, cytokeratin 5 and nuclei respectively in *Adgra3^cre/cre^* mice. Control *Adgra3+/+* in S1A. **I)** X-gal stained whole mounts of P10 eyelids from *Adgra3^lz/lz^* mice showing meibomian glands (black ar-rowheads) devoid of Gpr125 and strong expression in eyelash follicles (red arrows) Scale bar 100µm. n=3.

Next, we submitted both strains of mice for evaluation by a veterinary ophthalmologist. Examination of the lens and retina by slit lamp revealed well-documented characteristics of control B6 and FVBN mice, but no abnormality specifically linked to the *Adgra3^cre/cre^* or *Adgra3^lz/lz^* genotypes. Fluorescein staining revealed no evidence for corneal abrasion, but highlighted the presence of large mucoid precipitates around the eyelids of homozygous mutants (Figure 2D). This feature pointed towards abnormal tear film composition. Tears are required to lubricate corneal and conjunctival surfaces and to prevent eyes from desiccation (Botelho, 1964). They also function to protect eyes from microbial infection and preserve visual acuity. Tear film is composed of three layers, each secreted from a different source (Figure 2E). Goblet cells, clustered along the conjunctival rim, provide the inner mucus layer that spreads tear film evenly over the ocular surface (Gipson, 2016; Rios et al., 2000). Meibomian glands, found between eyelash follicles on the inner surface of eyelids, produce the outer lipid layer that prevents evaporation (Bron and Tiffany, 1998; Nien et al., 2010). Lacrimal glands secrete the central aqueous component that contains water-soluble immuno-active and antibacterial proteins, as well as glucose, urea, and salts (Makarenkova et al., 2000). Defects in the volume or composition of any layer destabilizes tear film and induces DED prompting us to investigate the tear glands in more detail (Pflugfelder and de Paiva, 2017; Schaumberg et al., 2003; Schaumberg et al., 2002). As *Adgra3* mutant mice had swollen eyelids we looked first for changes in goblet cells and meibomian glands. Histological sections of eyelids stained with Alcian blue revealed goblet cells in *Adgra3^cre/cre^* mice (Figure 2F) but with greater variation in number (average =70/eyelid; range 7-210 n=23) compared to controls (average of 65 goblet cells/eyelid; range 45-94; n=23): some showed epithelial and goblet cell desquamation next to swathes of mucus; others showed clusters of goblet cells. Meibomian glands displayed inflammatory infiltration by T-cells and macrophages (Figure 2G,H). Surprisingly, given these phenotypes, goblet cells and meibomian glands were devoid of Gpr125 expression whereas eyelash follicles were positive in X-gal stained eyelids (Figure 2I). These data show that changes in goblet and meibomian glands seen in human DED occur of *Adgra3* mutants (Pflugfelder and de Paiva, 2017; Schaumberg et al., 2003; Schaumberg et al., 2002) but as a secondary consequence of a tear film abnormality caused by loss of Gpr125 elsewhere. By a process of elimination this led us to focus on the lacrimal gland.

### *Adgra3^cre/cre^* and *Adgra3^lz/lz^* mice have abnormal lacrimation

We tested whether Gpr125 loss affected lacrimal function by measuring tear volume. *Adgra3^cre/cre^* and *Adgra3^lz/lz^* mice produced two to three-fold more tears than heterozygous or wild-type controls (Figure 3A). Tear volume was greater in female than male mice. *Adgra3^cre/cre^* and *Adgra3^lz/lz^* mice often presented with a mild phenotype in one eye (squint only) and a severe phenotype (blepharedema and or mucus) in the other. When we separated eyes into mild and severe categories according to photographic assignment taken prior to measurement and then reanalyzed the data, we found tear volume for the mild phenotypic category was similar, and sometimes lower, than those of wildtypes. In contrast, those in the severe category showed high values indicative of excessive tearing (Figure 3A). Thus, our mice recapitulated the paradoxical phenomenon documented in human patients with DED where individuals with tear film abnormality originating from initial mild ocular dryness respond to the consequent eye irritation with compensatory hyper-lacrimation (Pflugfelder and de Paiva, 2017).

**Figure 3.**
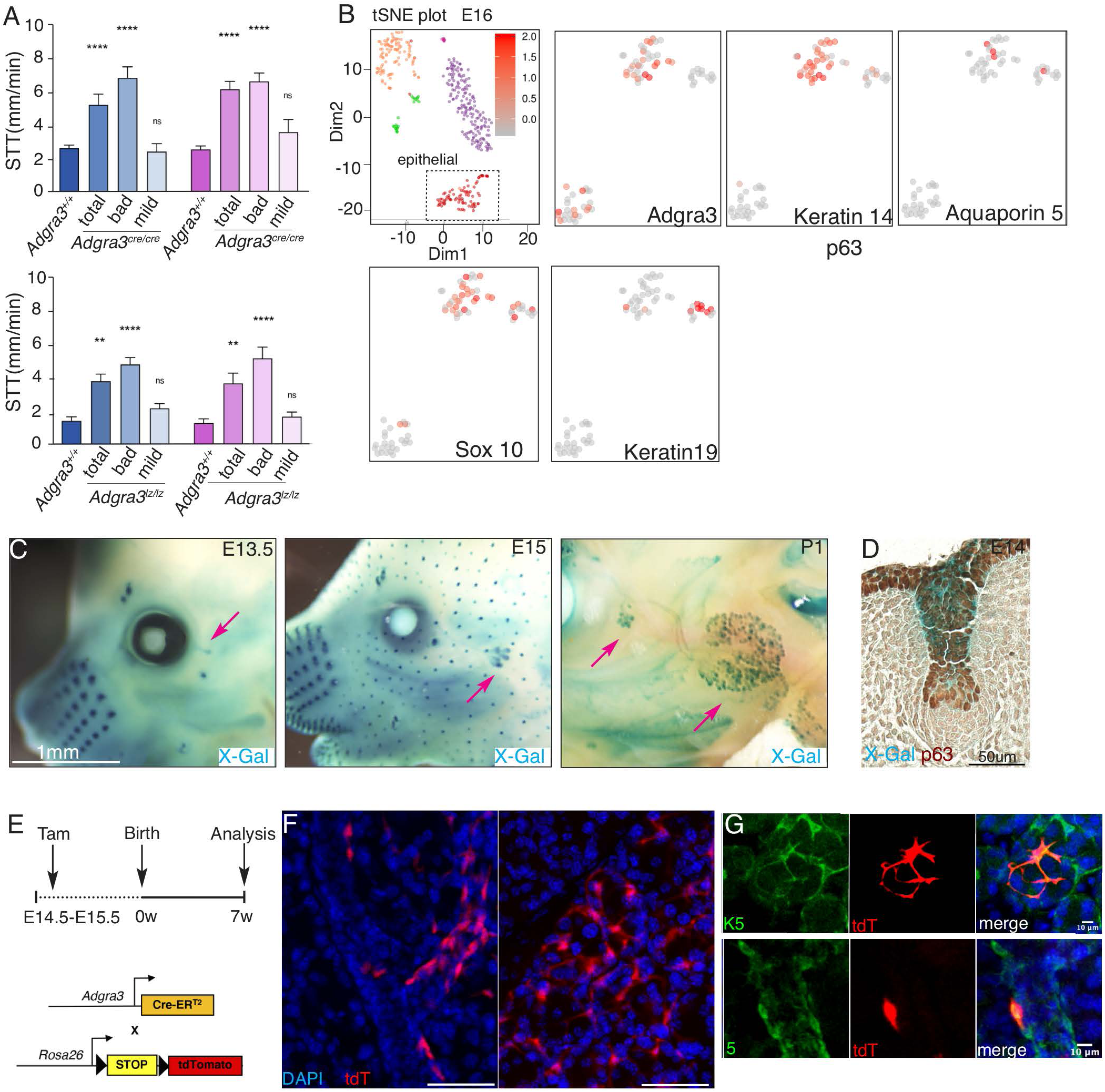
Gpr125 cells, located at ductal tips during development, function as lacrimal myoepithelial progenitors. **(A)** Increased tear production observed in *Adgra3^cre/cre^* and *Adgra3^lz/lz^* male (blue) and female (pink) mice compared to controls. Each bar represents the mean ± SEM on 6-32 mice. **** p<0.0001, **, p<0.05 value significant; ns, not significant. **B)** t-SNE plot of cells clusters within E16 lacrimal glands (10). Zoomed images of E16 epithelial compartment (boxed region) show cells expressing Adgra3 mRNA also express myoepithelial markers, Keratin14 and Sox10 but not luminal markers Keratin19 or Aquaporin5. **C)** Gpr125-β-gal expression in embryos and in the bulge **(D)** of E14 whisker follicle stained with p63. **E)** Strategy for tracing the lineage of Gpr125-positive cells in E14.5-E15.5 embryos carrying the *Rosa26.*lox.STOP.lox.TdTomato reporter by tamoxifen injection of pregnant *Adgra3^cre/cre^* dams. 3D-confocal images of lacrimal glands from mice at **(F)** 7 weeks and **(G)** 6 months showing tdT expression in elongated myoepithelial cells along the basal border of ducts and stellate cells enmeshing acini colocalized with myoepithelial marker (K5). Scale bar 50µm.n=3.

### Gpr125-expressing cells are located at the leading tips of ducts during lacrimal development and function as progenitors of the lacrimal myoepithelium

Given the significant effect of *Adgra3* loss on lacrimal function, we sought to identify cell types that express Gpr125 over the course of lacrimal development. We began by mining scRNAseq data (Farmer et al., 2017). Gpr125 mRNA was detected in a small cell population that co-expressed keratin 14 and Sox10 mRNAs (Figure 3B). This population was present during the early developmental stage of ductal elongation (E16) but diminished by P4 (data not shown) as acinar differentiation ensued. Next, we stained embryos with X-gal to locate cells expressing Gpr125 (Figure 3C). Murine lacrimal glands emerge around embryonic day 13 (E13) as a bulbous outgrowth of the conjunctival epithelium, which by E15 has elongated as a bi-layered hollow duct with 4-5 bulbous tips.

Subsequent branching produces a compact mass of secretory acini, enmeshed by contractile myoepithelial cells, which fully differentiate after birth (Dean et al., 2004; Dean et al., 2005; Farmer et al., 2017; Makarenkova et al., 2000). Gpr125-β-gal appeared in the lacrimal bud as it emerged from the conjunctival rim ∼E14, but by E15.5 it was restricted to a discrete population of cells located at the leading tips of migrating lacrimal ducts and by P1 at the front of lacrimal branches (Figure 3C). A potential role in directional outgrowth and collective cell migration is suggested by the fact that Gpr125 contains LRIG motifs that are present in Slit/Robo guidance factors (Bjarnadottir et al., 2006; Simundza and Cowin, 2013). This concept is supported by studies in zebrafish have shown that Gpr125 expression levels impact upon the migration of facial branchiomotor neurons (Li et al., 2013). Gpr124, is required for tip cell function during angiogenic sprouting raising the possibility that this family of proteins may serve similar roles in distinct cell types (Anderson et al., 2011; Vanhollebeke et al., 2015).

Intriguingly, during the course of these experiments we noted that Gpr125-β-gal was also expressed within a well-characterized “bulge” stem cell compartment of hair follicles and whiskers (Figure 3C, D) (Cotsarelis et al., 1990). This prompted us to ask if the embryonic Gpr125-positive cells present at ductal tips functioned as lacrimal progenitors. To test this, we performed lineage tracing by using the *creER^T2^* cassette present in *Adgra3cre* mice to activate expression of a lineage reporter. We labeled embryos harboring the ROSA-lox-STOP-lox-tdTomato reporters at E13-E15 by delivering tamoxifen to *Adgra3^cre/cre^* dams during mid-pregnancy. Lacrimal glands were harvested at 7 weeks and 6 months of age and analyzed by 3-D immunofluorescence confocal microscopy (Figure 3E,F). At 7 weeks, we found tdTomato (tdT)-labeled cells with an elongated shape along the basal borders of ducts and with stellate morphology enmeshing acini. These characteristics, together with their expression of keratin 5 (K5), identified them as contractile myoepithelial cells. A similar pattern was seen in glands from mice harvested at 6 months, indicating significant longevity of the original progenitor population (Figures 3F,G). Collectively, these data show that Gpr125+ cells at tips of migrating embryonic lacrimal ducts function as long-lived unipotent progenitors of the ductal and acinar myoepithelium. Recent studies have traced K14, K5, Sma and Runx lineages in the lacrimal gland and revealed a complex hierarchy comprising multipotent Runx cells at the apex and lineage restricted progenitors arising before birth (Basova et al., 2020). Our study adds to these analyses by providing a highly specific cell surface marker of Sma+ Sox10+ unipotent myoepithelial progenitors, locating these progenitors at the distal tips of elongating embryonic ducts, and demonstrating that they are already lineage restricted between E13-E15 of embryonic lacrimal development. Our findings of Gpr125 in stem cell compartment of several tissues together with early documentation of its expression in spermatogonial progenitors, show Gpr125 has widespread utility as marker for the localization and isolation of early progenitors.

### *Adgra3^cre/cre^* and *Adgra3^lz/lz^* mice show inflammatory infiltration of lacrimal glands

Given Gpr125 expression in myoepithelial progenitors and its homology to adhesion receptors, we investigated the effect of Gpr125 loss on myoepithelial integrity of lacrimal glands. In histological sections we noted the presence of foci, composed of small round cells where the lacrimal acinar organization was disrupted in *Adgra3^cre/cre^* (Figure 4A,B) and *Adgra3^lz/lz^* mice. Immunostaining for K5 was conspicuously absent in these foci indicating that myoepithelial cells were lost or disrupted (Figure 4C,D). These foci were surrounded by F480-positive macrophages (Figures 4C,E), and filled with cells recognized by CD4 and CD8 antibodies (Figures 4C, F,G) indicating infiltration by T-helper cells and cytoxic T-cells. Histological analysis of females displaying facial swelling (Figure 4H) and proptosis during pregnancy and lactation revealed enlarged lacrimal glands (Figure 4I,J) with swathes of macrophages around large areas of acinar loss (Figure 4K). Thus, loss of Gpr125 is accompanied by impaired lacrimal myoepithelial integrity and lymphocytic infiltration. Inflammatory infiltration is a common feature in later stages of human DED and a central feature of Sjogren’s syndrome, the third most common auto-immune disease (Pflugfelder et al., 2018; Pflugfelder and de Paiva, 2017; Schaumberg et al., 2003; Schaumberg et al., 2002).

**Figure 4.**
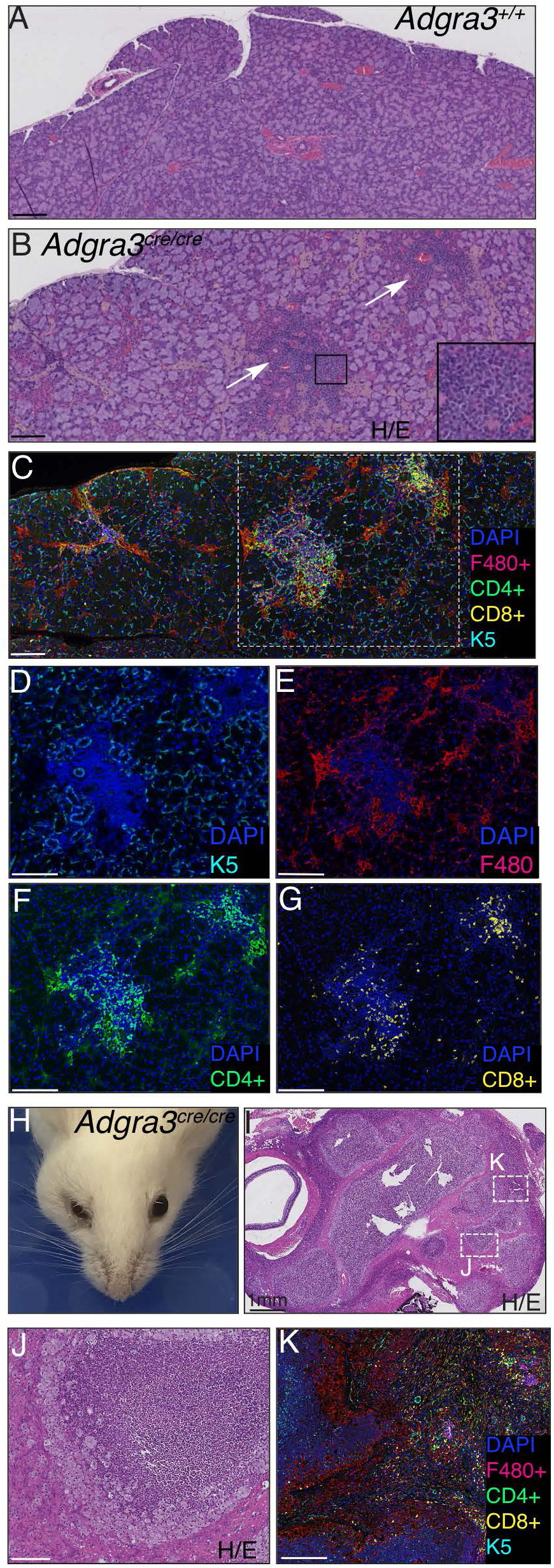
Loss of Gpr125 leads to abnormal lacrimation and inflammatory infiltration of the lacrimal glands. **(A)** H/E section of lacrimal gland from *Adgra3^cre/cre^* mice shows foci of infiltration (arrows). **(B)** Control *Adgra3+/+*. **(C-G)** Immunofluorescence of lacrimal gland co-stained for **(D)** K5, **(E)** macrophages, **(F)** T-helper, **(G)** cytotoxic T cells. **(H)** *Adgra3^cre/cre^* female with lacrimal mass sectioned and stained with H/E in **(I,J)**. **(K)** Immunofluorescence analysis as described above of boxed region in **I**. Scale bar 100µm.

Analysis of knockout (KO) mice has revealed a suite of critical regulators of tear film (Chen et al., 2014; Cui et al., 2005; Dean et al., 2004; Dean et al., 2005; Gipson, 2016; Kenchegowda et al., 2011; Makarenkova et al., 2000; Marko et al., 2013; McMahon et al., 2014; Plikus et al., 2004; Tong and Gupta, 2016; Tsau et al., 2011). However, the involvement of GPCRs in this process has not been studied. In sporadic DED, tear film abnormality exposes the eye to irritation and desiccation, prompting compensatory excessive tearing and immune response, which leads to further lacrimal destruction (Pflugfelder and de Paiva, 2017). Our *Adgra3* mutants reproduce this complex spectrum of symptoms, from early eye discomfort to blepharedema (Figure 1), mucus accumulation (Figures 1, 2D), goblet cells desquamation (Figure 2F), compensatory hyper-lacrimation, and inflammatory infiltration of meibomian (Figures 2H) and lacrimal glands (Figure 4B-G). Moreover, *Adgra3* mutants show worsening of their eye phenotype during pregnancy and lactation, recapitulating the hormonal/gender epidemiology of DED, which is more prevalent in women and exacerbated by pregnancy and post-menopause (Schaumberg et al., 2003; Schaumberg et al., 2002). Many genetic mouse models of DED arise from immune dysregulation or defects in matrix inhibition of immune activation and thus recapitulate late stages of DED (Park et al., 2015; Tong and Gupta, 2016). In contrast, our mice identify an initiating event during lacrimal development that predisposes mice to the full pathophysiological progression of DED. We show that in the absence of Gpr125, focal areas of the lacrimal gland become devoid of myoepithelium (Figure 4D) and infiltrated by lymphocytes and macrophages (Figure 4C-G). These results reinforce the concept that myoepithelial cells play a critical role in DED and complement studies that have shown that myoepithelial differentiation and contractile function is altered in Sjogren’s patients (Hawley et al., 2018; Makarenkova and Dartt, 2015). Going forward, it will be important to determine if Gpr125 is involved in blepharitis and DED in humans. DED is a significant health problem that affects ∼5% of the population overall is particularly prevalent in the elderly and women (Pflugfelder and de Paiva, 2017; Schaumberg et al., 2003; Schaumberg et al., 2002). As GPCRs are currently the targets of approximately 34% of drugs approved by the US Food and Drug Administration then deciphering Gpr125 signaling pathways holds promise to uncover novel targets for therapeutic intervention in these common conditions (Bassilana et al., 2019).

## Materials and Methods

### Mice

Mice were constructed by Ingenious Technologies, Ronkonkoma, NY as follows. A cassette containing *CreER^T2^* followed by a 3’ polyadenylation signal, harboring SV40- driven Neo flanked by FRT sites inserted in a central intron, was recombined into a bacterial artificial chromosome (BAC) to place *CreER^T2^* under the control of the *Adgra3* promoter, excising 502 bp encompassing 221 bp of exon 1 and part of the following intron 1-2 of *Adgra3*. Mice generated from these ES cells were selected for germline transmission by PCR, verified by southern analysis and sequencing then bred to a Flp deleter strain to remove Neo. *Adgra3^lz/+^* mice were generated by Regeneron using VelociGene methods to modify a bacterial artificial chromosome (BAC) clone carrying the mouse *Adgra3* gene by replacement of sequence encompassing exons 16-19 with *lacZ* to produce expression of fusion protein comprising the N-terminal extracellular domain, the first transmembrane domain, and part of the first intracellular loop of Gpr125 fused to β-galactosidase (Figure 1A) (Seandel et al., 2007; Valenzuela et al., 2003). Animal experiments were approved by NYUMC institutional animal care and use committee and conformed to American Association for Accreditation of Laboratory Animal Care guidelines.

### Ophthalmologic examination

Standard ophthalmic examination was performed by a trained veterinary ophthalmology consultant (Dr. Michael Brown, Animal Eyes of New Jersey). Slit lamp biomicroscopy was used to assess the cornea, anterior chamber, iris, lens, and vitreous humor. Mydriasis was induced with tropicamide and the retina was examined via indirect ophthalmoscopy. Corneal fluorescein staining was performed by applying sodium fluorescein (1%), for 3 minutes to the cornea of mice. Excess fluorescein was removed by flushing with sterile phosphate buffered saline (PBS) and corneal staining was evaluated and photographed with a slit lamp biomicroscope (Humphrey-Zeiss, Dublin, CA) using a cobalt blue light. Punctate staining was recorded using a standardized National Eye Institute grading system of 0 to 3 for each of the five areas of the cornea.

### Schirmer Tear Test

Tear production was measured via a modified Schirmer Tear Test. Briefly, 35mm x 5mm wide commercial Schirmer Tear Test standardized sterile strips (Schirmer Tear Test; Merck Animal Health) were transected with into two 15mm x 2.5mm strips, with the top notch removed. Individual strips were placed under the lower eyelid using forceps and removed after 15 seconds. The length of dye migration and wetting of the strip was measured in millimeters under a dissecting microscope.

### Intraocular pressure measurement

Mice were anesthetized and maintained on isoflurane through a nose cone. IOPs were measured using a TonoLab rebound tonometer (Icare, Finland) within 5 min after isoflurane gas anesthesia induction. For every 6 valid measurements, the highest and lowest IOP values were automatically excluded by the device, and the average of the remaining 4 IOP values was displayed along with the deviation. For quality control, only averages with slight deviation of less than 2.5 mmHg were considered acceptable readings. This procedure was repeated at least 3 times for each eye, and the acceptable readings were averaged IOP was measured eighteen times for each eye, and the average value was used for final analysis.

### Histological Analysis

The exorbital lacrimal gland, salivary and parotid glands, and whole globes were removed from mice and fixed with either 10% neutral buffered formalin or 4% paraformaldehyde (PFA) and embedded in paraffin. For general histological assessment, sections were stained with hematoxylin and eosin (H&E), or with periodic acid-Schiff (PAS) and Alcian Blue to visualize conjunctival goblet cells. Goblet cells in the bulbar and palpebral conjunctiva were quantified by two separate readers. Serial sections of tissues were stained with antibodies for anti-CD4, anti-CD8, anti-F480, anti-cytokeratin CK5 optimized by the Experimental Histology Core, NYUMC for analysis by Akoya/PerkinElmer Vectra® multispectral imaging system then counterstained with Dapi.

### Lineage Tracing

For lineage tracing experiments, *Adgra3*^creERT2^ mice were crossed to the fluorescent Rosa26R-lox.STOP.lox-tdTomato lineage reporter strain (Stock No. 007909) Jackson laboratory. The transcriptional STOP was deleted by cre recombination during embryonic development (E14.5-E15.5) by delivering tamoxifen (5mg per mouse-2 doses of 2.5mg) by oral gavage to *Adgar3*^^cre/cre^^ dams with during mid-pregnancy. Pups were delivered at E19.5-E20.5 by caesarian section to avoid problems with delivery caused by Tamoxifen and fostered by SWR/J mice. Their tissues were harvested at 7 weeks and at 6 months to test for progenitor potency and longevity.

### Tissue clearing and 3-D imaging

Lacrimal glands were excised and fixed overnight in 4% PFA then processed using a modified CUBIC (Reagent 1A) protocol (Davis et al., 2016). Tissue was incubated in CUBIC Reagent 1A clearing solution for 4 days, rinsed 3X in PBS then immunostained for 4 days at 4C in PBST containing 10% rabbit serum and rabbit anti-K5 (Covance, PRB160P, 1:100), rinsed again then 2 days in goat anti-rabbit AlexaFluor (AF) 647 (Thermo Fisher Scientific, A21245, lot number 1805235, 1:500), rinsed 3X then cleared in CUBIC R2 for 24hrs. Cleared lacrimal tissues were imaged using a Zeiss 880 Laser Scanning inverted confocal microscope with 20X air Plan-Apochromat N.A. 0.8 M27 objective lenses.

### X-gal staining

Embryos, Eyes and lacrimal glands were fixed in 4% paraformaldehyde (PFA) (Sigma-Aldrich) at room temperature (RT) for 30-60 min, rinsed 3X in X-gal rinse buffer (2 mM MgCl2, 0.1% Sodium deoxycholate, and 0.2% NP-40 in PBS) at RT, then incubated in X-gal staining solution (50 mg/ml 5-bromo-4-chloro-3-indolyl-β-D-galactopyranoside in rinse buffer containing 5 mM potassium ferricyanide, 5 mM potassium ferrocyanide) (Applichem, Cheshire, CT) at RT overnight. After staining, glands were rinsed in PBS, post-fixed in 4% PFA overnight then prepared for whole mount analysis or processed for paraffin embedding and sectioned for histological analysis.

### scRNA-seq analysis

scRNA-seq analysis were generated mining data from Farmer DT. et al. (Farmer et al., 2017) We processed the dataset using iCellR, Single (i) Cell R package, an interactive R package to work with high-throughput single cell sequencing technologies with the help of NYU Langone’s Applied Bioinformatics Laboratories (https://www.biorxiv.org/content/10.1101/2020.03.31.019109v1).

### Statistical Analysis

Experimental data are presented as mean ± SEM. P values for experiments comparing two groups were calculated using student’s t test. For experiments comparing more than two groups, an Ordinary one-way ANOVA was used with multiple comparisons test. P<0.05 was considered statistically significant.

## Online supplemental material

Fig. S1 (related to Fig. 2) shows immunostained meibomian gland in control *Adgra3+/+ versus Adgra3^cre/cre^* mice.

## Acknowledgements

This work was supported by Department of Defense W81XWH-17-1-0013 BC160959 (PC), NIH training grants: T32GM066704 (JS), T32CA009161-44 (AI) and The Susan G Komen Foundation For The Cure (AI). We thank Applied Bioinformatics, Experimental Pathology Research and Microscopy Laboratory Cores for the service provided: Grants P30CA016087, S10 OD021747 and P30CA016087. We thank Dr. Aris Economides, Regeneron, Tarrytown NY for advice on *Adgra3-*creER^T2^ construction and Dr. Michael Brown DVM (Animal Eyes of New Jersey, NJ) for ophthalmological evaluation.

The authors declare that no conflict of interest exists.

Author Contributions: PC conceived the study, designed experiments and wrote the paper with input from ES and RB; ES RB JS AI MF and AS conducted the experiments.

---

**Figure S1.**
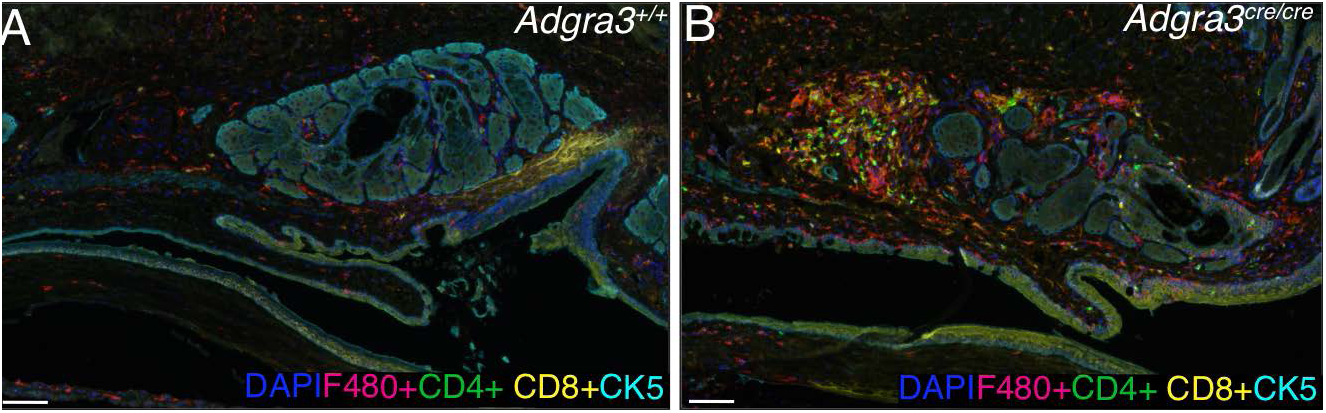
**(A,B)** Meibomian gland immunostained with antibodies: F480, CD4, CD8, CK5 and DAPI to detect macrophages, T-helper, cytotoxic T cells, cytokeratin 5 and nuclei respectively in control *Adgra3+/+* and *Adgra3^cre/cre^* mice. Scale bar 100µm.

## Notes

### Competing Interest Statement

The authors have declared no competing interest.

